# How the insect central complex could coordinate multimodal navigation

**DOI:** 10.1101/2021.08.18.456777

**Authors:** Xuelong Sun, Shigang Yue, Michael Mangan

## Abstract

The central complex of the insect midbrain is thought to coordinate insect guidance strategies. Computational models can account for specific behaviours but their applicability across sensory and task domains remains untested. Here we assess the capacity of our previous model explaining visual navigation to generalise to olfactory navigation and its coordination with other guidance in flies and ants. We show that fundamental to this capacity is the use of a biologically-realistic neural copy-and-shift mechanism that ensures sensory information is presented in a format compatible with the insect steering circuit regardless of its source. Moreover, the same mechanism is shown to transfer cues from unstable/egocentric to stable/geocentric frames of reference providing a first account of the mechanism by which foraging insects robustly recover from environmental disturbances. We propose that these circuits can be flexibly repurposed by different insect navigators to address their unique ecological needs.

## Introduction

Recently, it has been proposed that repertoire of robust navigation behaviours displayed by insects (***Webb and Wystrach, 2016***; ***Wehner, 2019***) can be traced to the well conserved brain region known as the central complex (CX) (***Honkanen et al., 2019***; ***Hulse et al., 2020***). The evidence to support this hypothesis includes: the discovery of the insect head-direction system in the CX that tracks the animal’s current heading relative to external (***Heinze, 2014***; ***Seelig and Jayaraman, 2015***; ***Kim et al., 2019***; ***Hardcastle et al., 2021***) or self-motion cues (***Green et al., 2017***; ***Turner-Evans et al., 2017***) cues; the innervation of the fan-shaped body (FB) region of the CX with sensory information relevant to different orientation strategies (***Hu et al., 2018***; ***Franconville et al., 2018***; ***Hulse et al., 2020***; ***Shiozaki et al., 2020***) (like sun compass (***Heinze et al., 2013***), vision (***Turner-Evans and Jayaraman, 2016***; ***Shiozaki et al., 2020***) etc.); the well-preserved columnar structure that is well suited to computing desired headings for vector navigation tasks (***Stone et al., 2017***; ***Honkanen et al., 2019***; ***Le Moël et al., 2019***; ***Lyu et al., 2020***); and the identification of a neural steering circuit in the FB capable of computing motor commands that reduce the offset between the current and desired headings computed (***Stone et al., 2017***; ***Honkanen et al., 2019***; ***Rayshubskiy, 2020***). Computational models of this architecture have produced realistic path integration (***Stone et al., 2017***; ***Gkanias et al., 2019***) and trap-lining behaviours (***Le Moël et al., 2019***), and simple conceptual extensions have been outlined that could generate long-distance migratory behaviour (***Honkanen et al., 2019***). Yet, for the CX to be considered a general navigation centre, three additional functions are required: generation of gradient ascent/descent behaviours that rely on spatially-varying but rotationally-invariant sensory cues (e.g. odour gradients); co-ordination of competing guidance systems into a single meaningful motor command; and demonstration of generalisation of function across sensory modalities and task spaces.

We recently demonstrated how the steering circuit could ascent gradients of visual familiarity when augmented by a neural *‘copy-and-shift’* mechanism that converts temporal changes in spatially sampled sensory information into an orientation signal (***Sun et al., 2020***). Specifically, the mechanism firstly *copies* the animal’s current heading from the head direction cells in the protocerebral bridge (PB) to desired heading networks in the FB. At the same time the signal undergoes a lateral *shift* in proportion to any undesired change in sensory valence as measured by the MB output neurons (***Aso et al., 2014***; ***Li et al., 2020***; ***Hulse et al., 2020***). Thus, the animal will continue on its current heading until an undesirable change in sensory valence is experienced at which point the shift mechanism will create an offset between the current and desired headings causing the steering circuit to initiate a change of direction. The architecture of the CX already possesses neural substrates ideally suited for both the *‘copy’* and *shift’* functions: heading direction cells are known to transmit their output into the ring structures of the central body (***Stone et al., 2017***; ***Honkanen et al., 2019***) as needed for *copy* stage; and neural mechanisms that laterally shift the head direction cells in response to sensory feedback (e.g. the self-motion cues (***Turner-Evans et al., 2017***; ***Green et al., 2017***), the visual cues (***Kim et al., 2019***; ***Fisher et al., 2019***)) are well established as required for the *shift* stage. Crucially, the complete *‘copy-and-shift’* mechanism explains how the CX steering circuit could exploit sensory gradients that provide no instantaneous orientation information for navigation.

We also demonstrated neural mechanisms that coordinate between different guidance strategies (***Sun et al., 2020***). Specifically we added a contextual-switching mechanism that triggers specific guidance strategies depending on the context, e.g. path integration unfamiliar contexts switching to visual route-following familiar contexts. As a final stage, we revealed how ring attractor circuits (***Touretzky, 2005***; ***Sun et al., 2018***) assumed to exist in the fan-shaped body provide an ideal substrate for optimally integrating cues that exist within a shared context (e.g. path integration and visual homing in unfamiliar contexts). The *‘copy-and-shift’* mechanism again plays a crucial role in this capacity as it “transfers” orientation outputs into a shared frame of reference. For example, when ascending gradients temporal changes in visual familiarity are translated into heading commands relative to the head-direction system which then share a frame of reference with the path integration system.

This biologically-realistic model of the insect midbrain was shown capable of generating realistic visual navigation behaviours of desert ants through the coordinated action of visual route following (RF), visual homing (VH) and path integration (PI) modules partially addressing two of the functional shortcomings listed above (***Sun et al., 2020***). In this study, we extend our analysis of the model, and in particular the *‘copy-and-shift’* mechanism, to assess if it can address the latter issue of generalisation across and between sensory and task domains. The following sections first assess whether the model can be easily reapplied to the olfactory tasks of chemotaxis and odour-gated anemotaxis (plume-following) in more laboratory olfactory navigation tasks. We then probe whether the same integration mechanisms can generalise to odour-gated switching in both flies and desert ants. Finally, we provide the first account of how the central complex could transfer orientation cues from an egocentric to a geocentric frame of reference which we propose can enhance the robustness of navigation.

## Results

### Core odour navigation behaviours using *copy-and-shift*

Here we assess the ease with which our visual navigation model generalises to olfactory navigation tasks.

#### Chemotaxis of odour gradients

Adult and larvae fruit-flies readily climb rewarding odour gradients by modulating their heading direction in direct response to the temporal change in odour concentration (***Gomez-Marin et al., 2010***; ***Nagel and Wilson, 2011***; ***Kim et al., 2011***; ***Schulze et al., 2015***; ***Jung et al., 2015***) mirroring our model’s approach to visual homing. Moreover, the neural pathways of olfactory processing are well established and only differ from our model in their sensory origins (antennal lobe (AL) to the lateral horn (LH) (***Gupta and Stopfer, 2012***; ***Roussel et al., 2014***) and mushroom bodies (MBs) (***Aso et al., 2014***; ***Hulse et al., 2020***)) before connecting to the CX through direct or indirect (hypothetically via superior medial protocerebrum (SMP) (***Plath et al., 2017***; ***Hulse et al., 2020***)) neural pathways. Thus by simply changing the input from optic to olfactory lobes (see ***Figure 1***A (left panel)) our model is able to adapt its heading to align with the positive odour gradient over successive steps (see ***Figure 1***B (left panel)). ***Figure 1***C (left panel) demonstrates the realistic chemotaxis behaviour generated by the model in a classic ‘volcano’ environment (***Jung et al., 2015***; ***Schulze et al., 2015***). ***Figure Supplement 1*** provides similarly realistic paths in other odour landscapes.

**Figure 1.**
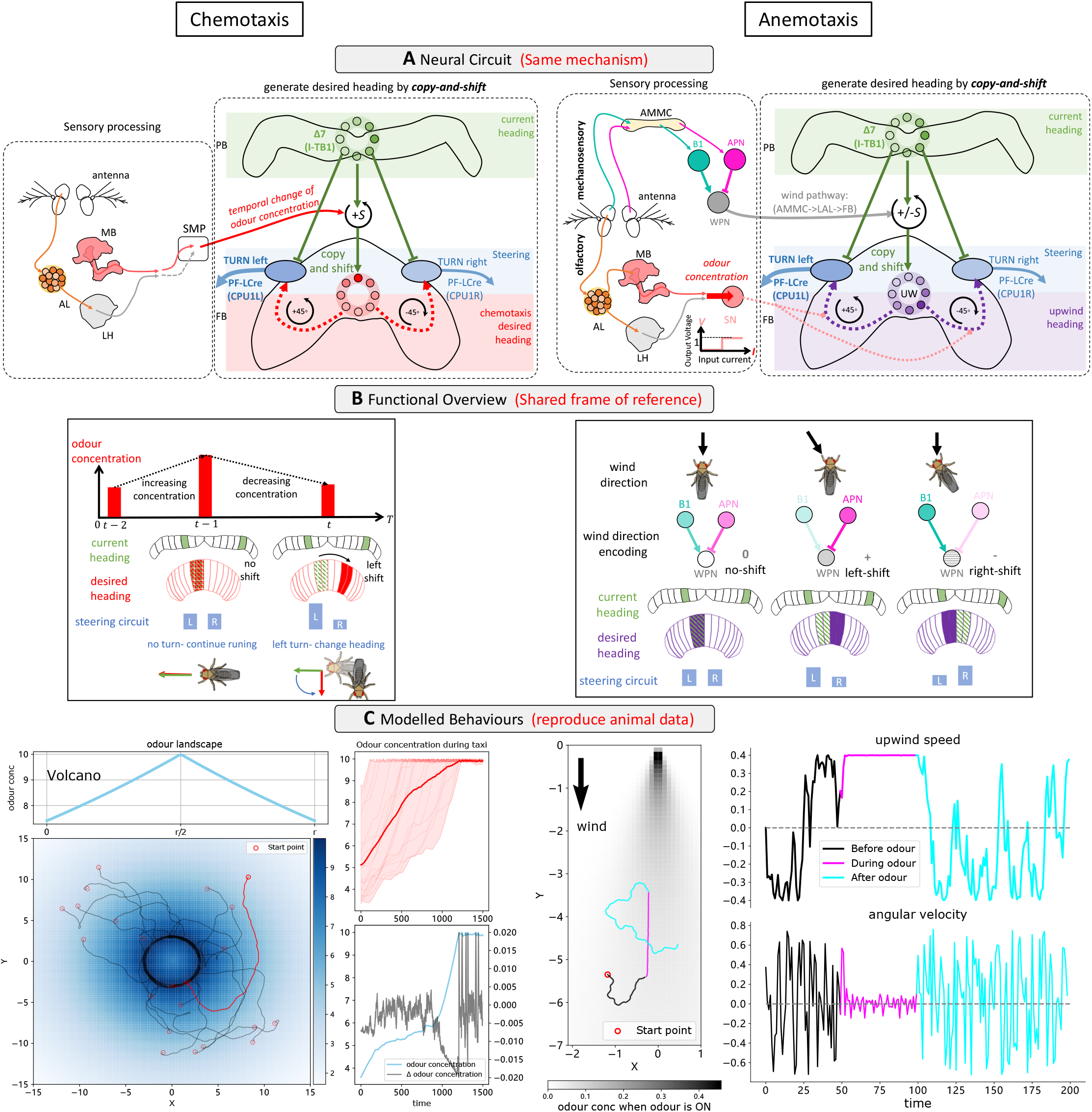
Modelling olfactory navigation in flies using ‘copy-and-shift’ mechanism: chemotaxis (left panel) and anemotaxis (right panel). **(A)**: Schematic diagrams of the neural circuits generating current-desired heading pairings for chemotaxis and anemotaxis. The same copy-and-shift mechanism is applied to ‘copy’ the same heading signal but with different driven signal for shifting, the temporal change of the odour concentration for chemotaxis and the wedge projection neuron (WPN) encoding the angular difference between the heading and upwind direction for anemotaxis. **(B)**: Schematic diagram of the functional explanation of the models. For chemotaxis, temporally decreasing odour concentration will shift the current heading to create the desired heading and then a turn can be made through the steering circuit. For anemotaxis, the WPN neuron subtracting the activation of the antennal mechanosensory and motor centre (AMMC) projection neuron (APN) from that of B1 will be the shifting source signal to generate the desired heading. Note that the two mechanisms share a frame of reference. **(C)**: The example behaviour generated by the proposed model mimicking the settings of the behavioural studies. For chemotaxis, simulated flies’ trajectories and perceived odour concentrations in a ‘Volcano’ odour landscape are shown. Increasing of the perceived odour concentration verifies the odour gradient ‘ascending’. For anemotaxis, magenta path segment is shown (when odour is ON) in compared with the undirected motion in the odour absence (when odour is OFF, black and cyan path segments). Upwind speed and angular velocity of the example agent are shown on the right panel. Note the obvious higher upwind translational velocity and low angular velocity during the presence of the odour indicating the ideal upwind surge manoeuvre. **Figure 1–Figure supplement 1.** The simulation results of chemotaxis model with odour landscape of ‘Linear’. **Figure 1–Figure supplement 2.** Simulation of wind direction encoding. **Figure 1–Figure supplement 3.** Simulation results of a group of agents (*N* = 20) driven by the odour-driven anemotaxis model.

#### Anemotaxis in odour plumes

In moving air-flows adult fruit flies pinpoint olfactory sources by anemotaxis whereby individuals align with the upwind direction allowing them to approach the hidden odour source (***Kennedy and Marsh, 1974***; ***Rutkowski et al., 2009***; ***van Breugel and Dickinson, 2014***). Insects sense wind direction through deflections of their antennae (***Yorozu et al., 2009***; ***Patella and Wilson, 2018***; ***Okubo et al., 2020***) which the wedge projection neurons (WPNs) convert into a direction relative to the animals current heading (***Suver et al., 2019***) (see ***Figure Supplement 2***). The WPN output is then transmitted to the FB of the CX via the antennal mechanosensory & motor centre (AMMC) -> lateral accessory lobe (LAL) -> noduli (NO) pathway (***Hulse et al., 2020***; ***Matheson et al., 2021***) (***Figure 1***B (right panel)). The *‘copy-and-shift’* mechanism again provides the ideal bridge between input signal and steering circuit. By simply driving the direction and magnitude of the *‘shift’* by the WPN response when a rewarding odour is detected (***Figure 1***A (right panel)) the model turns the agent upwind (see ***Figure 1***B (right panel)). ***Figure 1***C (right panel) shows an example path of a simulated fly navigating a classic laboratory environment with an odour plume into which rewarding odour is toggled ON and OFF (for a simulation of a group agents see ***Figure Supplement 3***), which demonstrates realistic odour-driven anemotaxis behaviour.

Taken together the above data demonstrates the capacity of the model to generalise from visual to olfactory navigation without significant alteration.

### Coordination of guidance behaviours by linking frames of reference

With the model shown to generalise from visual to olfactory navigation tasks, we now assess it’s ability to co-ordinate guidance strategies across sensory domains.

#### Contextual switching between olfactory guidance behaviours

In reality insects utlise both the chemotaxis and anemotaxis strategies outlined above. Across species and environments (laminar odour gradient or turbulent odour plume), a distinct behavioural trigger is reported at the onset (ON-response) or loss (OFF-response) of sensory valence (moths (***Kennedy and Marsh, 1974***; ***Rutkowski et al., 2009***), flying fruit flies (***van Breugel and Dickinson, 2014***), walking flies (***Gomez-Marin and Louis, 2012***; ***Steck et al., 2012***; ***Bell and Wilson, 2016***; ***Álvarez-Salvado et al., 2018***)). Specifically, in the presence of the attractive odour animals apply anemotaxis and surge upwind but when the attractive odour is lost they engage in a chemotactic-like search to recover the plume. This problem is analogous with the contextual switching using in our previous model to select between ON- and OFF-route navigation strategies reproducing the real ants data (***Wystrach et al., 2012***). ***Figure 2***A (left panel) depicts how the CX switching circuit can be easily reconfigured to be triggered by the instantaneous change of odour concentration fitting with the reported ON- and OFF-responses (***Álvarez-Salvado et al., 2018***). Note that we here assume that the ON- and OFF-response are driven by the output neurons of the odour processing brain regions (i.e., MBON or LHON) that could compute the temporal changes of odour concentration (***Hulse et al., 2020***; ***Matheson et al., 2021***). ***Figure 2***B (left panel) illustrates simulated ON- and OFF-responses that are supplied to the model and their behavioural consequence. ***Figure 2***C (left panel) demonstrates realistic olfactory navigation behaviour produced by the model which closely matches behavioural data in ***Álvarez-Salvado et al. (2018***). See also the simulation results of a 20-agents group demonstrating similar performance in ***Figure Supplement 1***.

**Figure 2.**
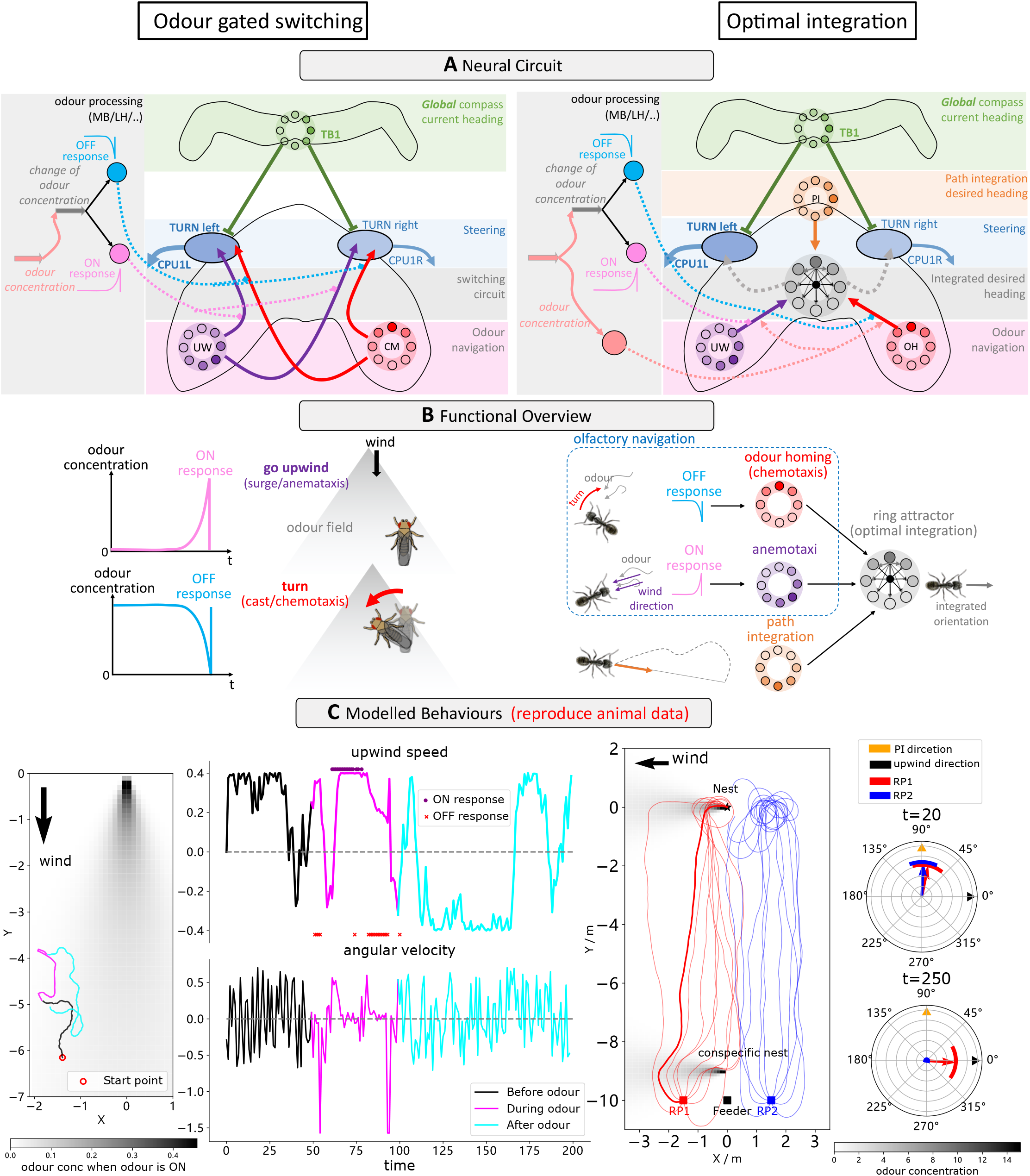
Neural models of optimally coordinating different desired headings sharing a same frame of reference. **(A)**: Schematic diagram sof the neural circuits. Left: temporal change of odour concentration based ON and OFF-response serving as the source signal of the switching circuit to select chemotaxis or anemotaxis strategy. Right: ring attractor network integrating multiple cues weighted by sensory values. **(B)**: Illustrations of the functional explanations of the model. Left: On-response triggers upwind following while OFF-response drive the animal to switch to chemotaxis leading a sharp turn which could bring the animal back into the odour plume. Right: ring attractor serves as the optimal integration circuit to integrate olfactory navigation including odour-driven anemotaxis and chemotaxis and the path integration. **(C)**: The example behaviour generated by the proposed model mimicking the real animal data. Left part of the left panel shows the trajectory of the one simulated fly, the upwind speed and angular velocity of the agent are shown in the right part. The time at which ON- and OFF- response is promoted also marked by purple star and red dots respectively. Left of the right panel draws the simulated ants’ homing paths and the group headings at *t* = 20 (early begin of the homing when PI dominates the steering) and *t* = 250 (nearly approaching the nest when olfactory navigation should dominate as PI vector length is low) are shown on the right. **Figure 2–Figure supplement 1.** The simulation results of a 20-agents group driven by the ON- and OFF-response based switching model. **Figure 2–Figure supplement 2.** Sensory perception and neural activities of the highlighted ant driven by the proposed model. **Figure 2–Figure supplement 3.** Simulation results where there is no conspecific nest near the releasing points with comparison to (C) right panel.

#### Optimally integration navigation behaviours across sensory domains

In barren salt-pans, homing desert ants follow their path integrator to their nest area before relying on nest-odour plumes for their final approach (***Buehlmann et al., 2012***). Ants bypass the nests of conspecfics that diffuse similar odours (*CO*_2_) until reaching the nest locale (***Buehlmann et al., 2012***) indicating use of a sophisticated integration strategy beyond simple switching outlined above. Rather, ants instead appear to weight their PI output relative to the home-vector length in a similar fashion to their integration of path integration and visual cues (***Wystrach et al., 2015***; ***Legge et al., 2014***) as was realised in our previous model using ring attractor networks (***Touretzky, 2005***; ***Sun et al., 2018, 2020***). ***Figure 2***A (right panel) depicts the augmentation of our odour-gated anemotaxis model with a ring attractor circuit to optimally integrate PI and olfactory navigation outputs. These adaptations are in accordance with the olfactory navigation mechanisms (chemotaxis and anemotaxis) proposed to be used by ants by ***Wolf and Wehner (2000***, 2005). Note that the desired headings recommended by odour homing (OH, or chemotaxis) and upwind direction (UW, or odour-gated anemotaxis) are gated by the OFF and ON response and weighted by the odour concentration signal prior to be injected to the RA to be combined with PI. ***Figure 2***B (right panel) illustrates how the various desired heading signals are optimally integrated by the ring attractor network prior to input to the steering circuit. ***Figure 2***C shows homing paths generated by the model following simulated displacements left or right of the regular feeder which closely match those of real ants (***Buehlmann et al., 2012***). Note that there is a additional odour plume diffused by a simulated conspecific nest positioned near the releasing points, for comparison, see the results of a simulation without this odour plume in ***Figure Supplement 3***.

Taken together these data demonstrate that the CX possess the neural mechanisms to flexibly coordinate the various guidance behaviours observed in insects across sensory domains supporting its role as the navigation centre (***Honkanen et al., 2019***; ***Hulse et al., 2020***).

### A mechanism for transferring between orientation frames of reference

The optimal integration model detailed above is reliant on the *copy-and-shift* mechanism firstly ensuring that all orientation cues are presented in a shared frame of reference. Recall that the desired headings for path integration, chemotaxis, and anemotaxis are all defined in relation to the animals’ global head direction. In the following analysis we assess whether this frame-changing capacity can also provide benefits for navigational robustness.

#### From egocentric wind direction to geocentric celestial compass

Desert ants travel to and from familiar feeder locations via visually guided routes (***Kohler and Wehner, 2005***; ***Mangan and Webb, 2012***) but wind gusts can blow them off course. ***Wystrach and Schwarz (2013***) reported that in the instant prior to displacement ants assume a stereotypical ‘clutching’ pose during which they transfer their egocentric measure of wind direction (indicating the direction in which they are about to be blown) into a geocentric frame of reference given by their celestial compass. Displaced ants then utilise this celestial compass memory to guide their path directly towards their familiar route (***Figure 3***A (left panel)). Such a strategy is easily accounted for by the *‘copy-and-shift’* mechanism as seen in ***Figure 3***B (left panel). That is, during the clutch pose the celestial compass heading is *copied*, and *shifted* by the activation of the WPN encoding the upwind direction relative to the animal’s heading to create a desired heading that points back along the direction of travel. This desired heading is maintained in a working memory during displacement before activation to guide the model back to the familiar route region (see simulated navigating paths in ***Figure 3***C (left panel)).

**Figure 3.**
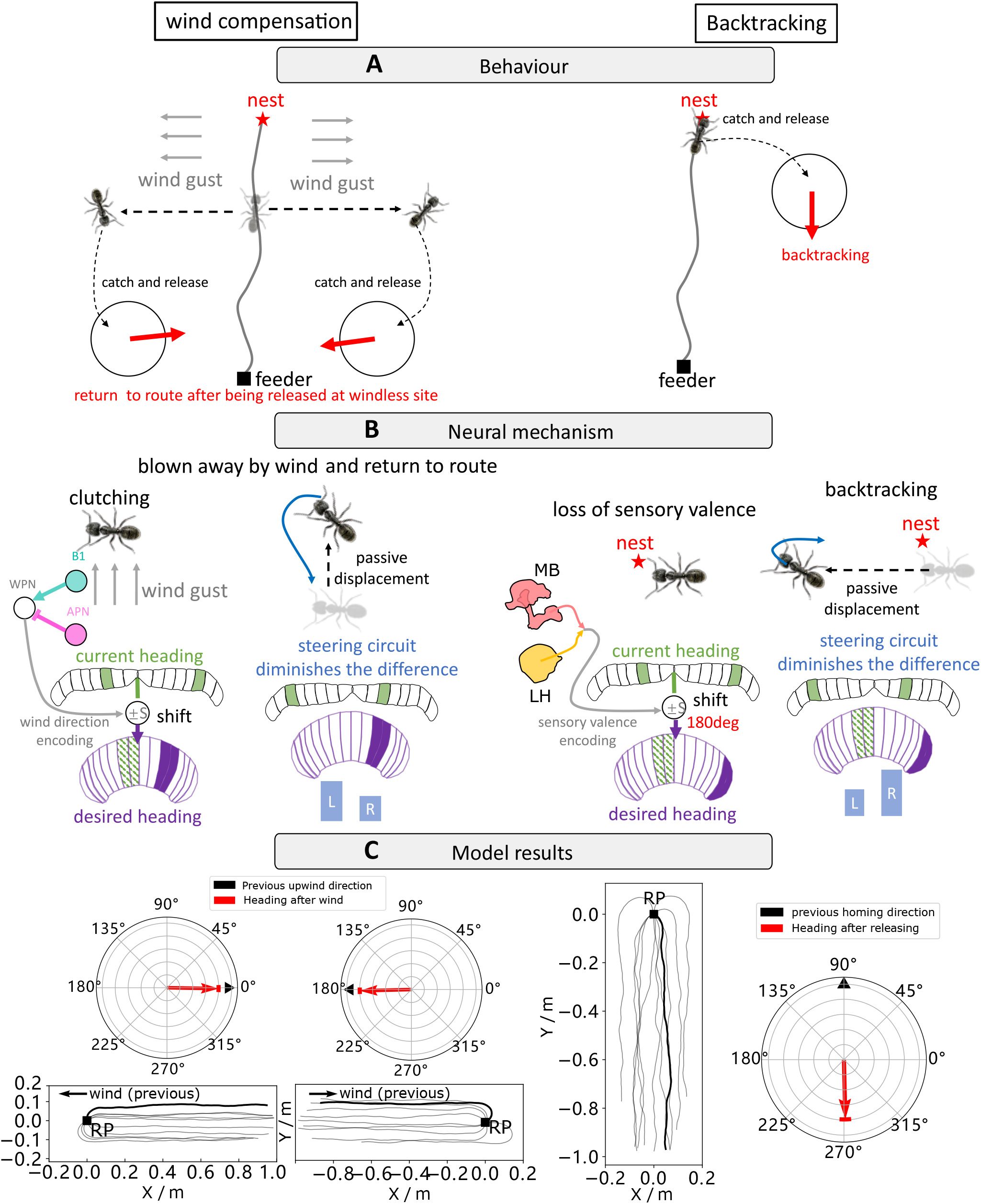
How the copy-and-shift mechanism endows insects with the ability to transfer egocentric frame of reference to the geocentric one for robust navigation. **(A)**: The wind compensation and backtracking behaviour of navigating ants. Left panel illustrates the wind compensation behaviour where the ants (with arbitrary initial headings) orientate to the direction of wind by which they have been blown away after been released in a windless site (***Wystrach and Schwarz, 2013***). Right subfigure shows that homing desert ants captured just before entering their nest and released in unfamiliar visual surroundings initially dash back along the celestial compass heading in which they were travelling (***Wystrach et al., 2013***). **(B)**: The proposed neural mechanism showing how the behaviours in (A) could be interpreted. Wind-compensation is implemented by using the *copy-and-shift* to *copy* their heading compass stored in the CX when clutching and *shift* by certain degree determined by the activation of WPN neurons to form the working memory (desired heading) for late navigation. Backtracking is modelled in identical way except that the *shift* is constant 180°. **(C)**: The simulation results of our model. In each panel, the navigating trajectories and initial headings of the simulated ants are shown. Simulated ants guided by the model are all heading to the expected orientation as observed in real behavioural experiments (***Wystrach and Schwarz, 2013***; ***Wystrach et al., 2013***).

#### From visual context to geocentric celestial compass

Similarly, homing desert ants captured just before entering their nest and released in unfamiliar visual surroundings initially dash back along the celestial compass heading in which they were travelling (***Wystrach et al., 2013***)(***Figure 3***A (right panel)). Note that this differs from the behaviour of ants lacking a path integration cues and displaced from other locations along the route who have no preferred direction of travel following displacement. This indicates that sight of the nest surroundings could be considered a ‘special circumstance’ in a similar way to the ‘clutching’ pose mentioned above. ***Figure 3***B (right panel) depicts how this behaviour could easily arise using the *‘copy-and-shift’* mechanism. That is, when there is a significant drop of visual novelty (as might only be experienced following displacement from the nest), the compass direction is again *copied* and *shifted* by a predetermined amount 180deg producing the ideal desired heading that can be stored in working memory and used to guide the initial stages of search as demonstrated in ***Figure 3***C (right panel).

In summary, the data above demonstrates the flexibility of the *‘copy-and-shift’* mechanism to transfer directional cues from an unstable frame of reference such as the wind direction to a stable frame of reference such as the global celestial compass. We proposed that this transfer is triggered by special sensory experience and motivational state of the animal, which provides another natural behaviour supported explanation of the numerous tangential inputs from multiple upstream brain regions to the FB (***Franconville et al., 2018***; ***Hulse et al., 2020***) forming a contextually dependent guidance network. This again extends the repertoire of guidance behaviour that the mechanism can account for and further supports to the role of the central complex as a navigation centre.

## Discussion

To summarise, we have shown how the CX-based steering circuit augmented with a *copy-and-shift* functionality can generate realistic odour-based chemotaxis and anemotaxis behaviours adding to the path integration, visual homing, visual route following, and long-range migrations explained previously (***Stone et al., 2017***; ***Honkanen et al., 2019***; ***Sun et al., 2020***). We have also outlined CX-based mechanisms that can coordinate guidance cues across sensory domains using biologically-realistic context-dependent switches and ring attractor networks. Finally, we demonstrated how the *copy-and-shift* mechanism can facilitate the transfer of orientation cues between unstable to stable frames of references. By triggering such a transfer under specific environmental conditions insects can increase the robustness of their guidance repertoire. The model presented can thus be considered as a general navigation model extending across multiple behavioural tasks (alignment with rotationally-varying compass, visual route or wind cues; and gradient ascent of spatially varying but rotationally-invariant cues such as odour and visual memories) experienced in multiple contexts. Taken together the results add further validation to the claim that the central complex acts as the seat of navigation coordination in insects.

The central complex is as ancient as insects themselves (***Homberg, 2008***; ***Strausfeld, 2009***) and is highly conserved across different species solving different navigational tasks (***Honkanen et al., 2019***; ***Hulse et al., 2020***). This fixed circuitry thus appears optimised to receive input from a variety of sensory sources and return a similar variety of navigational behaviours applicable across contexts. Indeed ***Doyle and Csete (2011***) posits that such ‘bowtie’ (or hourglass) architectures are also observed in the decision making circuits of the mammalian brain (***Redgrave et al., 1999***; ***Humphries and Prescott, 2010***) and function by providing *“constraints that deconstrain”* (see ***Figure 4***A). That is, the fixed circuitry of the CX constrains the format of the sensory input but decontrains the application domains of the output behaviours. Through interpreting various navigation behaviours by applying the same *‘copy-and-shift’* mechanism, our model demonstrates a specific implementation of such bowtie structure within the CX (***Figure 4***B).

**Figure 4.**
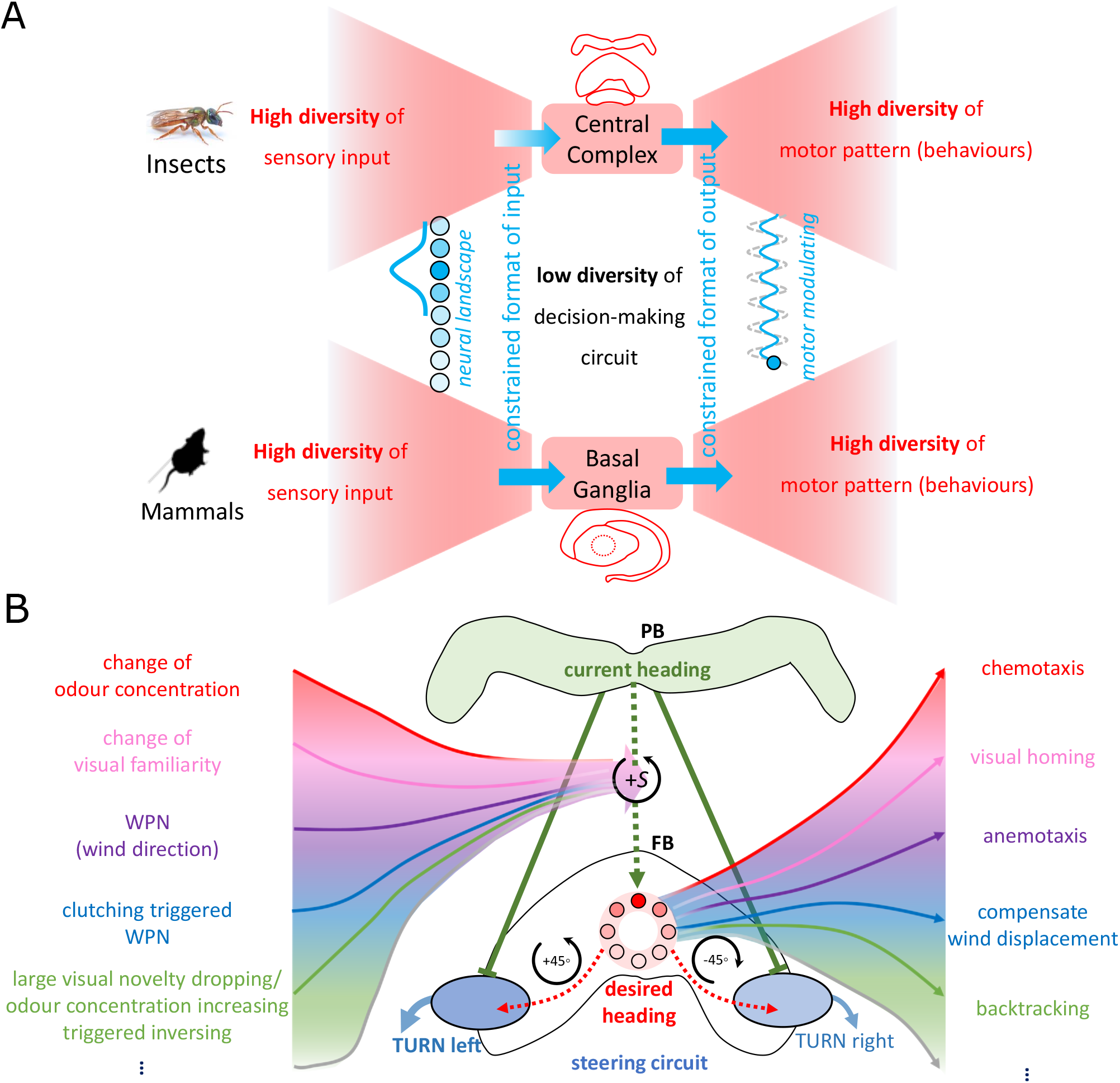
The ‘bowtie/hourglass’ architecture (*Doyle and Csete, 2011*) of biological control system. (A) The control system of insect navigation (top) and mammalian decision-making (bottom) are epitomised by the ‘bowtie’ architecture, proposing that fixed brain circuitry constrains the format of the sensory input (fanning in to the knot) but decontrains the application domains of the output behaviours (fanning out of the bowtie). (B) The proposed specific implementation of the bowtie architecture in modelling the CX for insect navigation. Specially, the copy-and-shift mechanism (regarded as the knot of the bowtie thus constrains the representation) that can be generally used to generate different desired headings across sensory and task domains (deconstrains the motor pattern thus allows for high diversity of behaviours).

This study has explored the behavioural consequences of the mechanisms using abstracted neural implementations, raising the question as whether they can be realised in insect brains. Regarding the *copy-and-shift* mechanism, lateralised neural connections and synapse-plasticity that shift the head-direction output relative to sensory input (i.e. nudge the activation ‘bump’ within a population of neurons) have already been mapped (***Seelig and Jayaraman, 2015***; ***Green et al., 2017***; ***Kim et al., 2019***; ***Fisher et al., 2019***) and modelled (***Cope et al., 2017***) demonstrating the feasibility of such computation. More recently, ***Goulard et al. (2021***) presented a CX-based navigation model that includes a biologically realistic neural pathway that is functionally similar to the *copy- and-shift* mechanism proposed here. The same study also outlined how a short-term memory of a desired heading could be maintained in the FB of the CX via synapse-weight modulation after the original guidance cue is removed, supporting the wind-compensation and backtracking behaviours described above. Lastly, our model hypothesises the existence of a ring attractor network to combines desired heading cues that mirrors the structure of the ring attractor in the head-direction circuit. The recently revealed 2D-grid backbone that is consists of vΔ (vertical) and hΔ (horizontal) neurons inter-connecting FB column neurons offers a potential substrate for such a network (***Hulse et al., 2020***), but additional investigations are required to verify this possibility.

Despite growing agreement on the functional role of the CX in insect navigation (***Honkanen et al., 2019***; ***Hulse et al., 2020***), a number of issues remain. Firstly, as well as innervating the CX, both visual and olfactory cues are also transferred directly to motor centres (***Rayshubskiy, 2020***; ***Scaplen et al., 2021***; ***Green et al., 2019***) providing redundant information streams. One possibility is that the direct pathways are used for fast reflex-like movements, whereas the CX pathway is responsible for higher-level guidance that requires learning and integration of multiple elemental guidance systems (***Currier et al., 2020***; ***Matheson et al., 2021***). This view is consistent with ***Steinbeck et al. (2020***) who demonstrate that the lateral-accessory-lobes (LAL), downstream of the CX, possess neural structures well suited to integrating outputs of the fast and the slow pathways. Future work is needed to merge these concepts into a single computational framework. Secondly, insects possess a MB in each brain hemisphere posing the question as to their combined role. ***Le Möel and Wystrach (2020***); ***Wystrach et al. (2020***) offer the hypothesis that MBs form an opponent memory system that can drive visual route following by balancing the difference in their outputs. This approach can be easily extended to incorporate both attractive and repulsive MB output neurons extending the application space and robustness of navigation. Integration of dual MB inputs represents an obvious next extension of the model presented here. Finally, the model presented here is unique in the format of the sensory data input to the MBs, and the behavioural strategies that the MBs generate. Specifically, we propose that the MBs process rotationally-invariant but spatially-varying cues (e.g. odour and visual familiarity gradients) and are thus responsible for generating gradient ascent/descent behaviours such as visual homing and chemotaxis via operant connections to the CX. In contrast, all rotationally-varying cues (e.g. wind-direction, visual route memories, and celestial compass) innervate the CX directly via alternate pathways (e.g. LAL). This separation of sensory information is fundamental to the flexibility of the model presented to create the array of behaviours presented and offers a testable hypothesis for future work. Such insights will be invaluable for refinement of our understanding of the robust navigation behaviours facilitated by the insect minibrain.

## Methods and Materials

All simulations and network models are implemented by Python 3.5 and external libraries-*numpy, matplotlib, scipy, opencv* etc. The source code of the simulation and plotting figures are available via Github.

### Odour field

As the basic sensory input, the spatial concentration distribution of the odour field is simulated simply and based on the scaled exponential functions, with required changes according to the wind dynamics.

#### Odour field without wind

For the simulations in the laminar odour environment (i.e. no wind) as that in ***Figure 1***(left panel), the landscape of the odour concentration *CON*_*o*_ are modelled for ‘volcano’ shape:

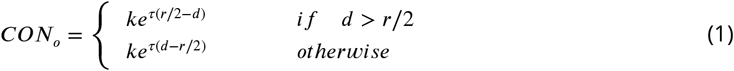

and for ‘linear’ shape:

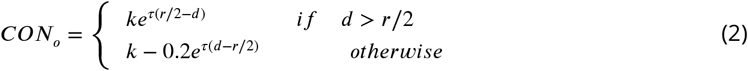

where *d* is the distance from the position (*x, y*) to the odour source (*x*_*s*_, *y*_*s*_). Thus, 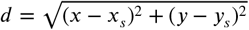 is the scale factor, *r* is the radius of the odour source and *τ* is decay factor.

#### Odour field with wind

To simplify the simulation of the odour plume dynamics, all the simulations in this study are conducted under the condition of constant wind speed *u* and wind direction *θ*_*w*_, and we assume that the odour plume will ideally flow to the downwind area, i.e., the odour concentration in the upwind area will always be zero. The source of the odour constantly emits at the rate *q*, Then the odour concentration at position (*x, y*) can be calculated by:

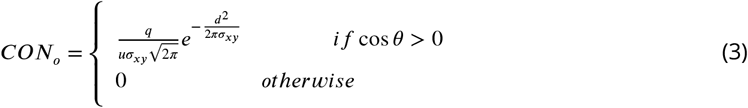

where 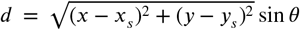 is the projected distance from the odour source. And *σ*_*xy*_ is calculated by *σ*_*xy*_ = *K*_*s*_*d* where *K*_*s*_ ∈ [0.5, 0.3, 0.2, 0.15, 0.1] is the tuning factor determined by the stability of the odour. And *θ* is the angel between the vector pointing from the position to the source and the wind direction, so can be computed by:

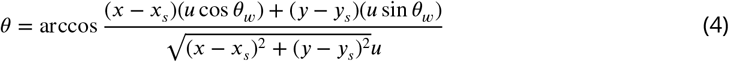

### Neural model

We use the simple firing rate to model the neurons in the proposed networks, where the output firing rate *C* is a sigmoid function of the input *I* if there is no special note. In the following descriptions and formulas, a subscript is used to represent the layers or name of the neuron while the superscript is used to represent the value at a specific time or with a specific index.

#### Current heading

In our previous model, there are two compass references derived from different sensory information (***Sun et al., 2020***), but in this paper, only the global compass, (i.e. the activation of I-TB1/Δ7 neuron) is used here because navigation behaviours reproduced in this study are all assumed using the global compass as the external direction reference. For the details of the modelling of global current heading 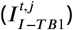 see our previous paper (***Sun et al., 2020***).

#### Steering circuit

The steering neurons (the same as previous paper (***Sun et al., 2020***) but presented here for convenience), i.e., CPU1 neurons 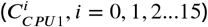 receive excitatory inputs from the desired heading 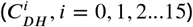 and inhibitory inputs from the current heading (*C*_*CH*_, *i* = 0, 1, 2…15) to generate the turning signal:

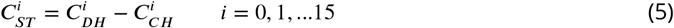

The turning angle is determined by the difference of the activation summations between left (*i* = 0, 1, 2…7) and right (*i* = 8, 9, 10…15) set of CPU1 neurons:

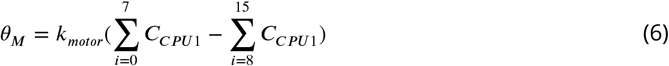

#### Upwind direction encoding

The upwind direction is decoded as the activation of UW neurons copied and shifted from heading neurons (I-TB1), the value of this shifting is determined by the angular difference between the current heading (*θ*_*h*_) and wind direction (*θ*_*w*_) encoded by the firing rate of WPN neuron. And the value of WPN neuron is defined as the difference of the antennal deflection encoded by B1 and APN neurons as:

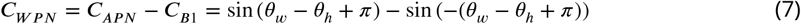

Then population activation of upwind direction neurons (UW) can be calculated by:

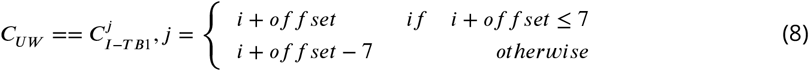

#### Fly-ON and OFF response based switching circuit

Different navigation strategy will dominate the motor system according to the sensory inputs, i.e., in this study, the change of perceived odour concentration. This coordination is modelled as a contextual switching that is very similar with the mechanism with SN1 and SN2 neuron involved in our previous model (***Sun et al., 2020***) to define the final output of odour navigation (*CO*_*N*_):

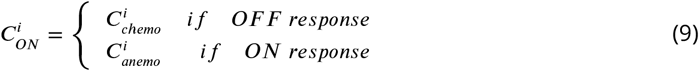

And how the sensory information determine the response is shown in ***Table 1***, where Random means no reliable sensory input is available, the agent will move forward to a random direction.

**Table 1.**
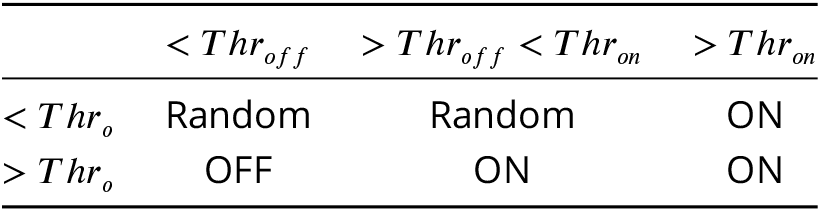
‘Truth table’ of the ON and OFF response of the modelled fly odour navigation. The column lists the state of sensed odour concentration while the row indicates the state of the changing of odour concentration.

##### OFF response-chemotaxis

The chemotaxis model is adapted from the previous visual homing model (***Sun et al., 2020***) by changing the change of visual familiarity signal from the MBON neuron (Δ*C*_*MBON*_) to the change of the odour concentration to determine the shifting value, thus the desired heading of chemotaxis is:

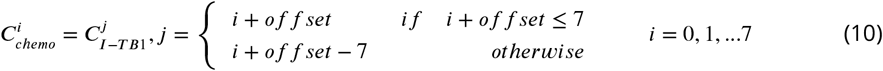

Note that, in our previous visual navigation model (***Sun et al., 2020***), *i, j* both are integer for the ease of computing, thus, the shifting accuracy is 45°, but here to more accurately model the desired heading and to achieve better performance, the shifting accuracy was set to be 4.5°by interpolating neuron activation of I-TB1 from 8 to 80 then down-sampling to 8 to generate shifted desired heading.

The relationship between the Δ*C*_*o*_ and the *of f set* is shown as following:

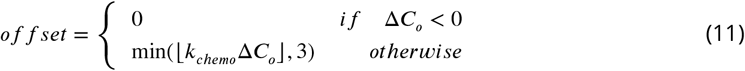

Then the desired heading of OH will be fed into the steering circuit to compare with the current heading to generate the motor command.

##### ON-response-dour-gated Anemotaxis

As shown in ***Table 1***, when the ON response is determined, the agent will follow the upwind direction, thus the desired heading input to steering circuit should be the upwind direction encoded by UM neuron ((8)):

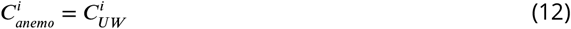

#### Ants-integration with PI

The modelling of ants’ odour navigation integrated with PI can be regarded as the extension of the fly’s odour navigation and an application of the unified model. Specifically, the final output of olfactory navigation is determined by the ON and OFF response (see ***Table 1***), and then is integrated with PI via RA like that in the optimal integration of PI and VH:

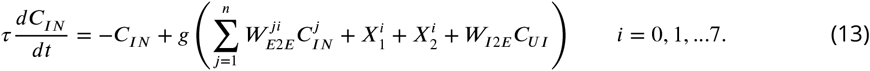

Where 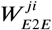 is the recurrent connections from *j*^*th*^ neuron to *i*^*th*^ neuron, *g*(*x*) is the activation function that provides the non-linear property of the neuron:

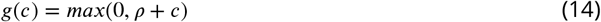

Where *ρ* denotes the offset of the function. Thus the *X*1 should be:

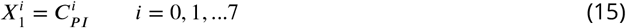

and *X*2 in (13) should be:

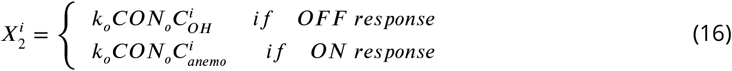

Then the output of optimal integration (OI) of the RA acts as the only desired heading input to the steering circuit:

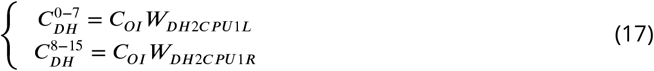

As only the global compass is needed in this study’s modelling. Thus the input of current heading will always be the excitation of the I-TB1 neuron:

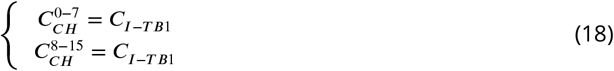

The output of the steering circuit (i.e., the summed activation of the left and right CPU1 neurons) is used to generate the turning command in the way that is same as (6).

### Simulations

In all simulations, at each time step, the simulated agent (walking fly or ant) will sense the odour sensory based on its current location and then update neural activation to generate the desired moving direction and finally move one step to that direction. ***Equation 6*** gives the turning angle of the agent, thus the instantaneous “velocity” (***v***) at every step can be computed by:

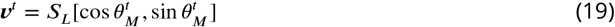

Where *S*_*L*_ is the step length with the unit of centimetres. Note that we haven’t defined the time accuracy for every step of the simulations, thus the unit of the velocity in this implementation is *cm*/*step* rather than *cm*/*s*. Then the position of agent ***P*** ^*t*+1^ in the Cartesian coordinates for the is updated by:

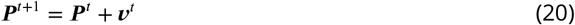

The position of odour sources in all simulations are all set to (0, 0), i.e., *x*_*s*_ = 0, *y*_*s*_ = 0. Other main parameters are listed in ***Table 2***

**Table 2.**
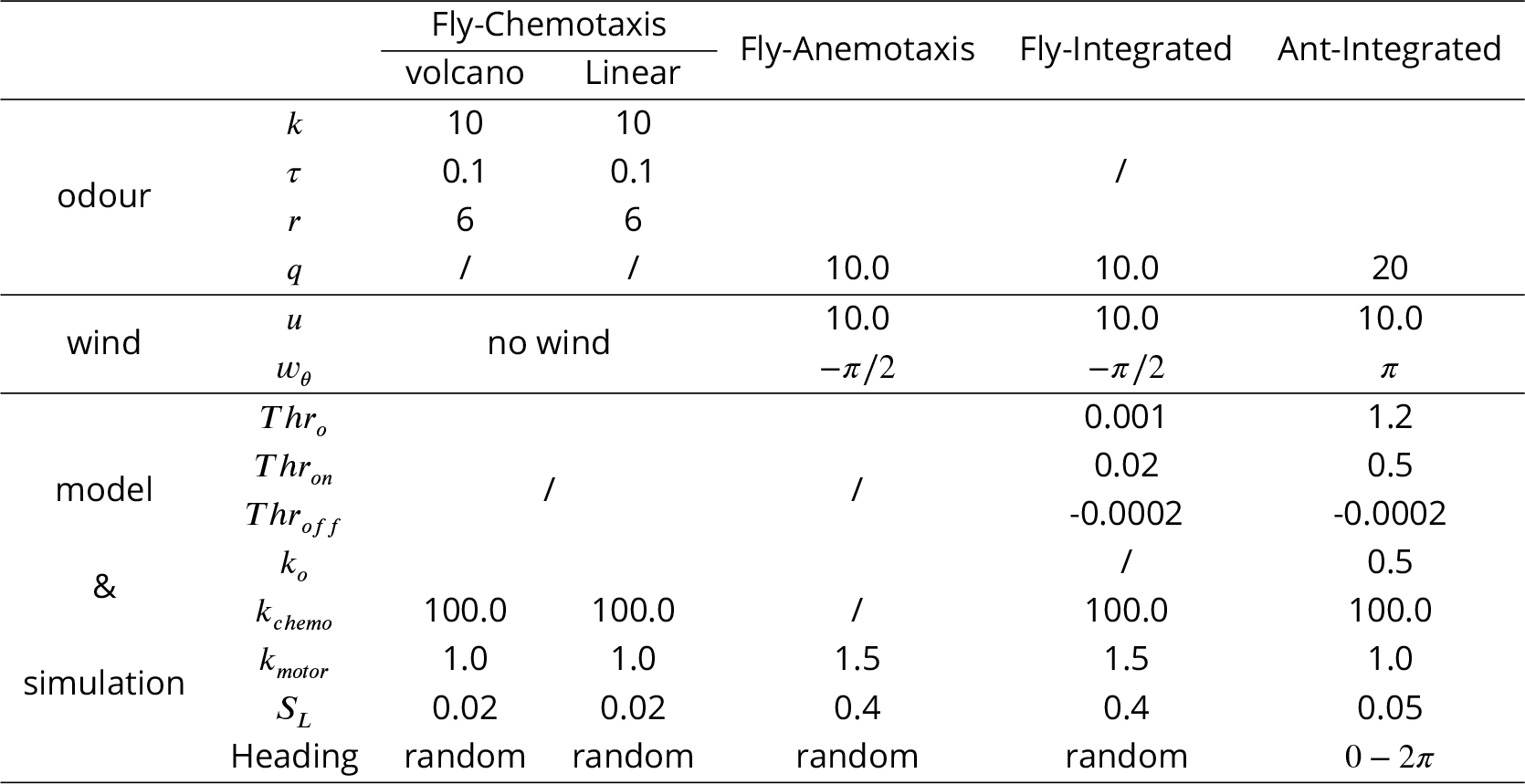
The detailed parameters settings for the simulations in this study.

#### Fly-Chemotaxis

To test the performances of the chemotaxis behaviour, 5 simulated agents with randomly generated heading direction starts from 5 randomly generated locations in the zone of (−12 < *x* < 12, −12 < *y* < 12), and then driven by the model for 1500 steps. Then we run this simulation for 4 times in two different odour landscapes (‘volcano’ and ‘linear’) to get the results shown in ***Figure 1*** (right panel) and ***Figure Supplement 1***.

#### Fly-Anemotaxis

To reproduce the behavioural data in ***Álvarez-Salvado et al. (2018***), the odour was only set on during the second a quarter of total time (e.g, if the agent is set to run 200 steps, then the odour- on time will in 50-100 steps). Four agents with randomly generated heading starts from randomly generated locations in the zone of (−1.5 < *x* < 1.5, −13 < *y* < −5), and then guided by the model to run 200 steps. The simulation was conducted for 5 times.

#### Fly-Integrated ON and OFF Response

The whole simulation settings are the same as that in the last section except for some model parameters listed in ***Table 2***, as this simulation is conducted to verify the integrated model.

#### Ants-Odour Navigation Integrated with PI

To reproduce the behavioural data in ***Buehlmann et al. (2012***), we first generate PI memory encoding the home vector with 10m length and *π*/2 direction. Then at each release point ((−1.5, −10) and (1.5, −10)), we released 10 simulated full-vector (10m-long and pointing to *π*/2) ants with different initial headings sampled uniformly from 0 − 2*π*, see also ***Table 2***. Note that the simulation settings with/without additional odour plume diffused by conspecific nest are identical so list as one column in ***Table 2***.

#### Antswind compensation and backtracking

The quick implementations of using ‘copy-and-shift’ mechanism to model the wind compensation and backtracking behaviour follow the same step: first, generate the desired headings by shifting the current heading by the WPN activation for the wind compensation and by 180°for backtracking respectively; second, release the simulated ant at the same releasing point but with random headings (uniform distribution in 0 − 2*π*). Motion-related parameters are set identically as that of Ants-Odour Navigation Integrated with PI.

## Acknowledgements

This research has received funding from the European Union’s Horizon 2020 research and innovation programme under the Marie Sklodowska-Curie grant agreement No 778062, ULTRACEPT and No 691154, STEP2DYNA.

---

**Figure 1–Figure supplement 1.**
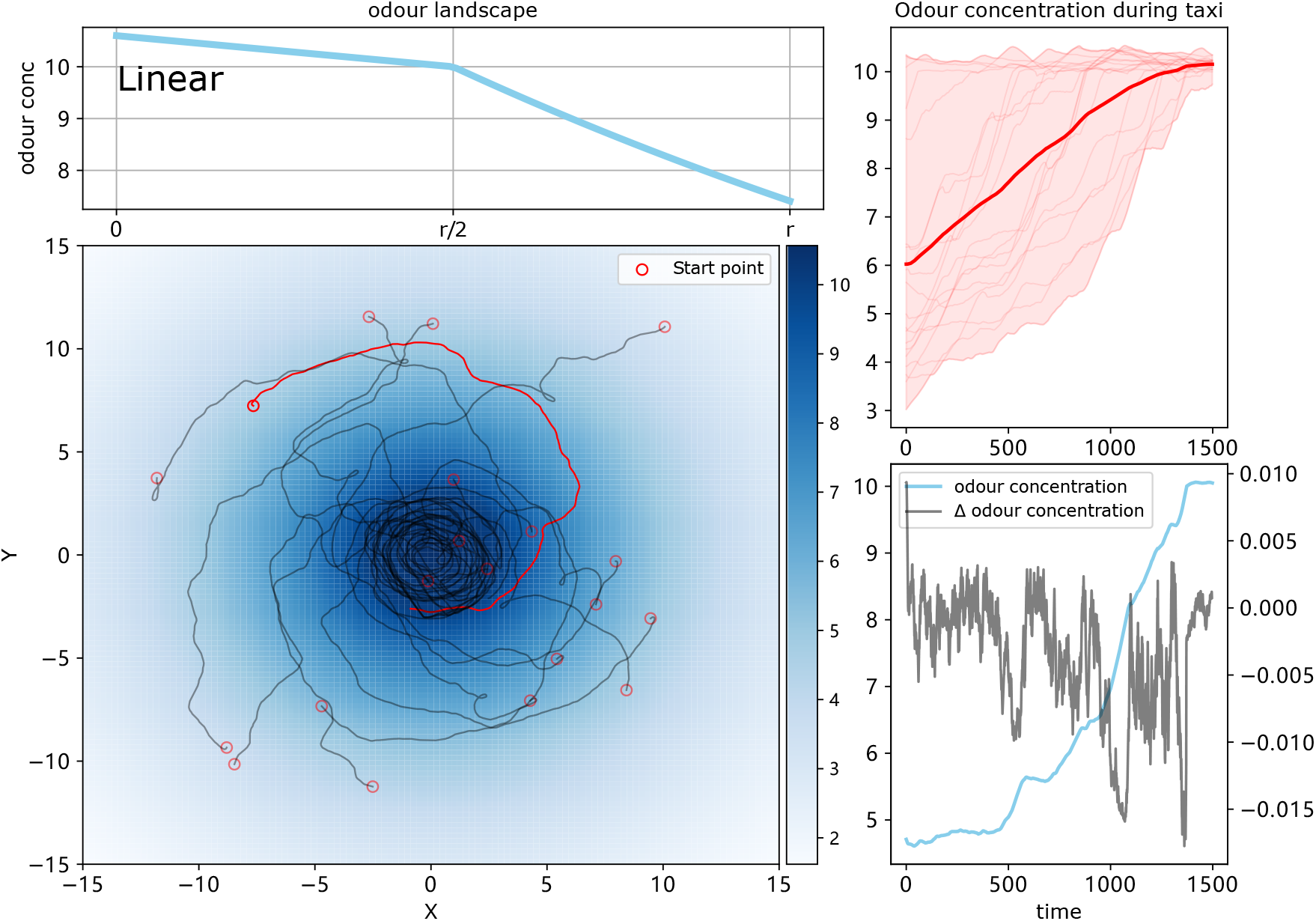
The simulation results of chemotaxis model with odour landscape of ‘Linear’. The odour field model and navigating trajectories are shown on the left whilst the perceived odour concentration and the temporal change of the highlighted agent are shown on the right.

**Figure 1–Figure supplement 2.**
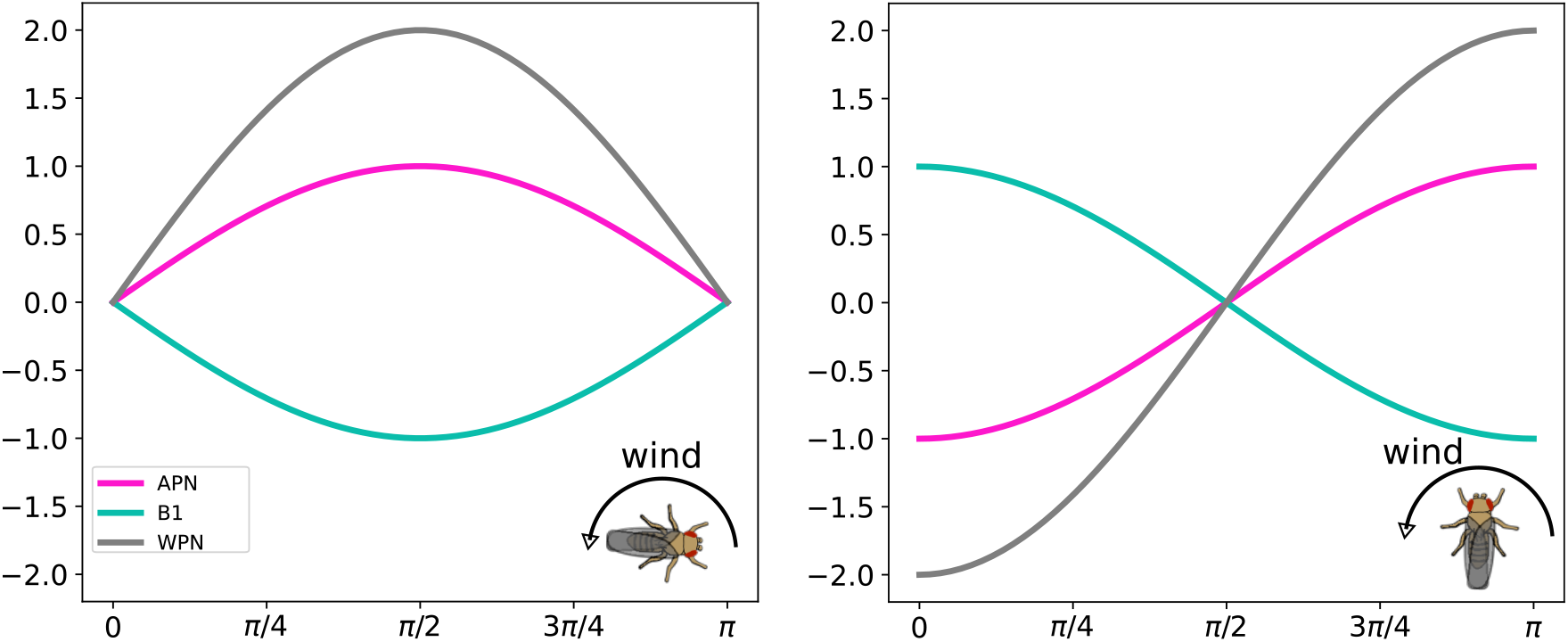
Neural responses of the wind direction encoding neurons with different animal headings (0 and *π*/2) and the wind direction stimuli is swept from 0 to *π*.

**Figure 1–Figure supplement 3.**
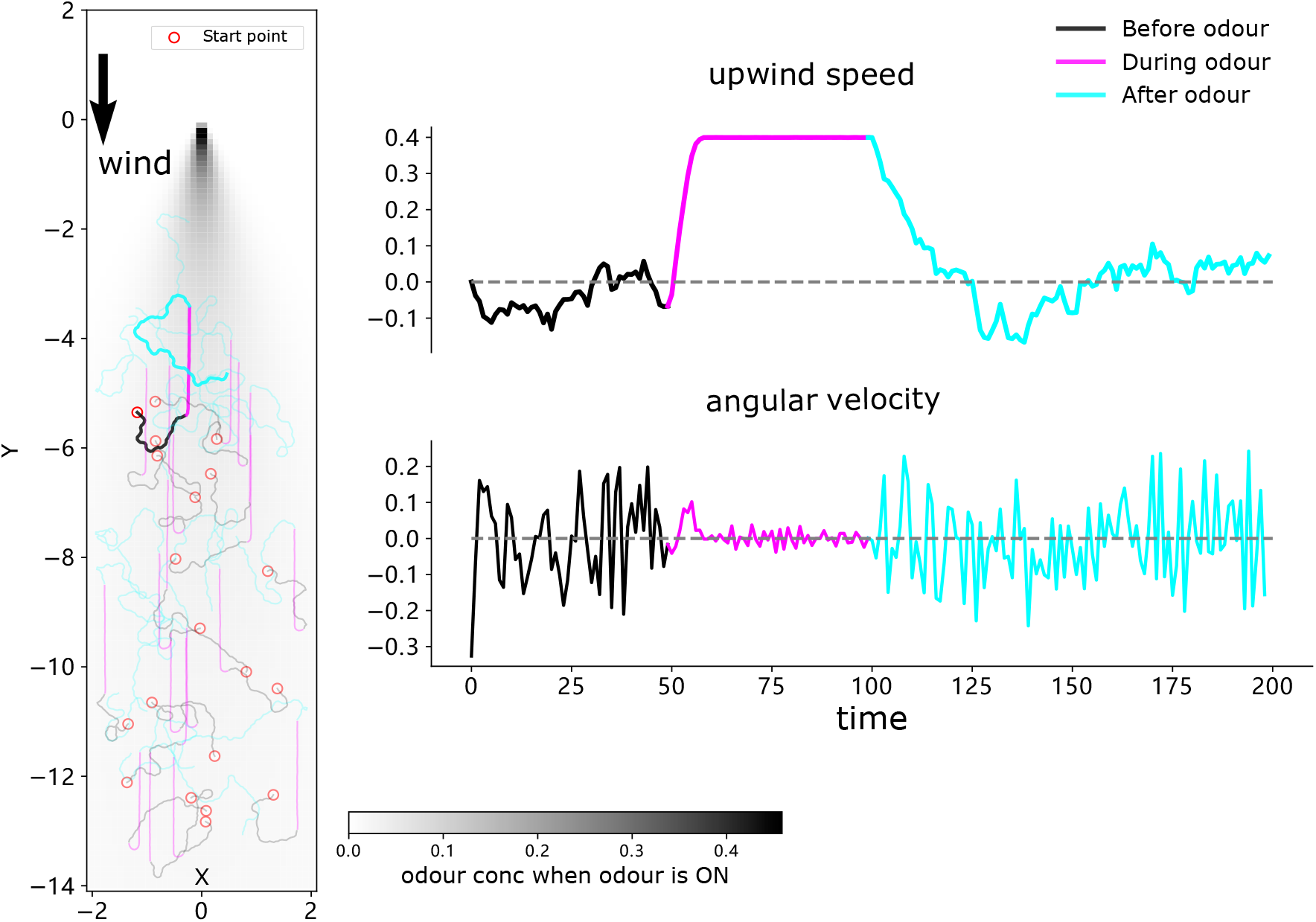
Trajectories of each agents (highlighted one corresponding to that shown in **Figure 1**), mean upwind speed and angular velocity of 20 simulated agents are shown.

**Figure 2–Figure supplement 1.**
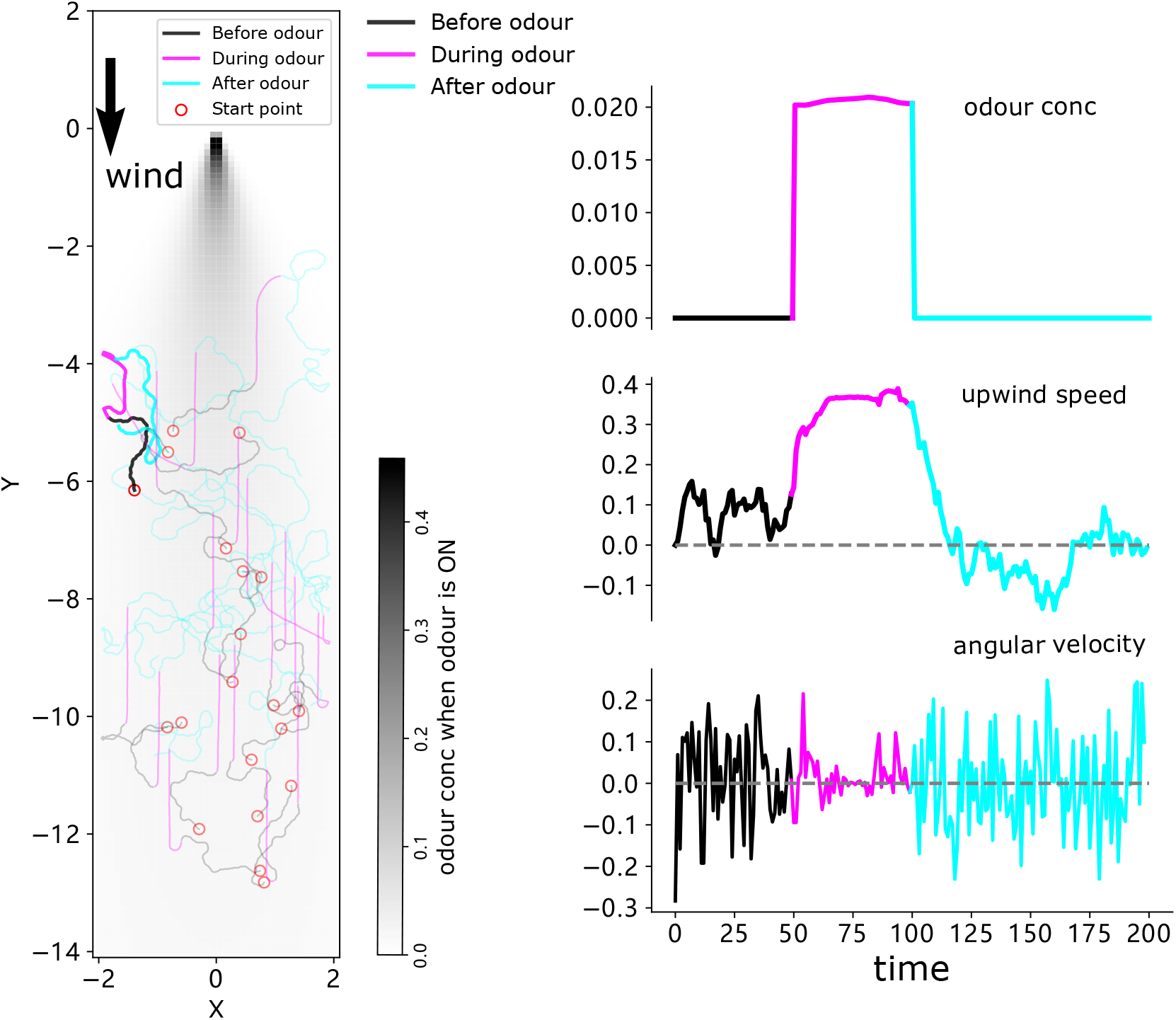
The simulation results of 20 agents. Trajectories are shown on the left with highlighted one corresponding to that of **Figure 2**, mean perceived odour concentration, upwind speed and angular velocity are plotted on the right.

**Figure 2–Figure supplement 2.**
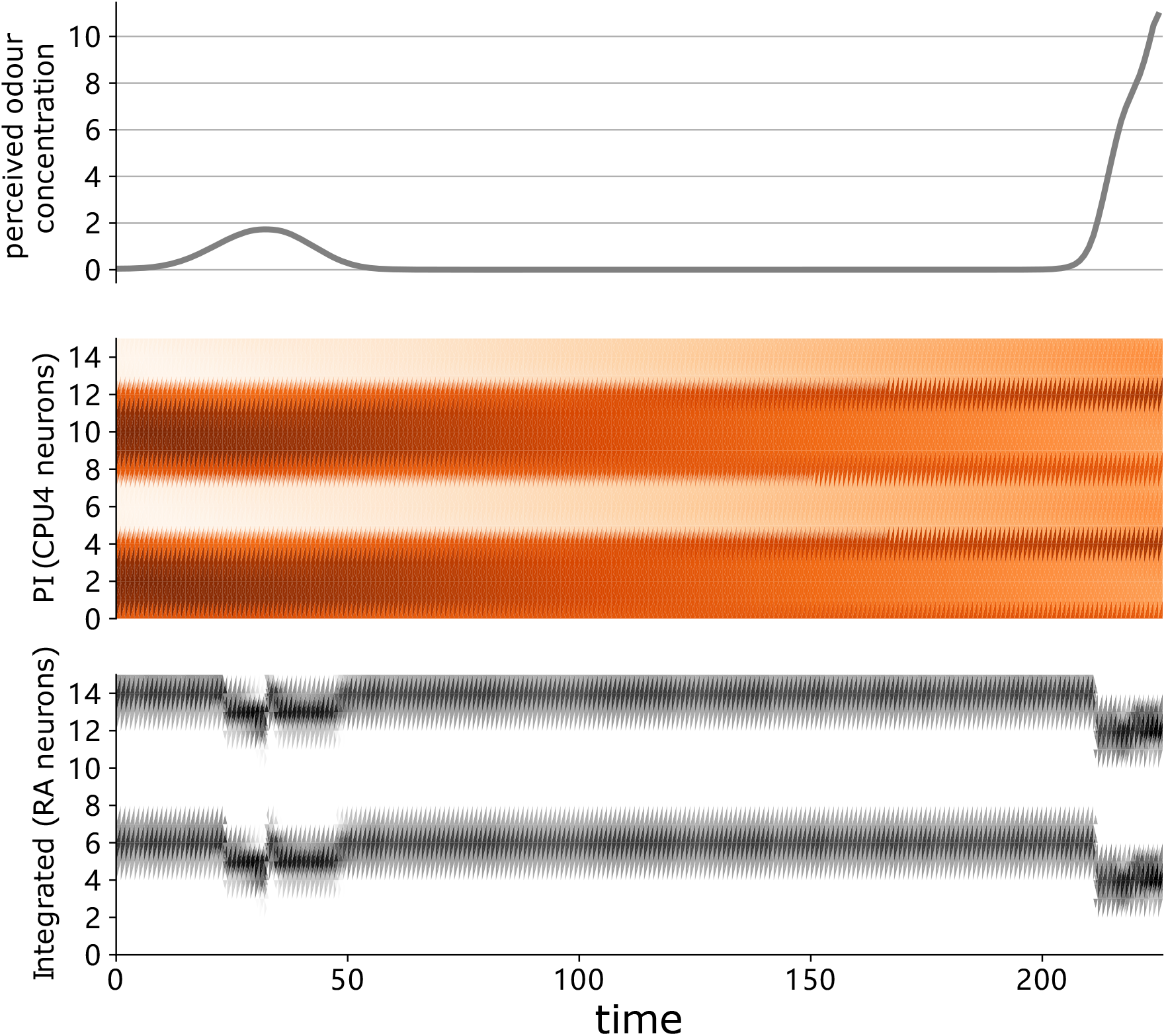
The instantaneous sensory value and neural activation of the highlighted agent in **Figure 2**C (right panel) during homing. From top to bottom, the value of perceived odour concentration, the activation of PI memory neurons (CPU4) and the ring attractor excitation neurons. Note that the output of the ring attractor neurons combines injected cues as expected.

**Figure 2–Figure supplement 3.**
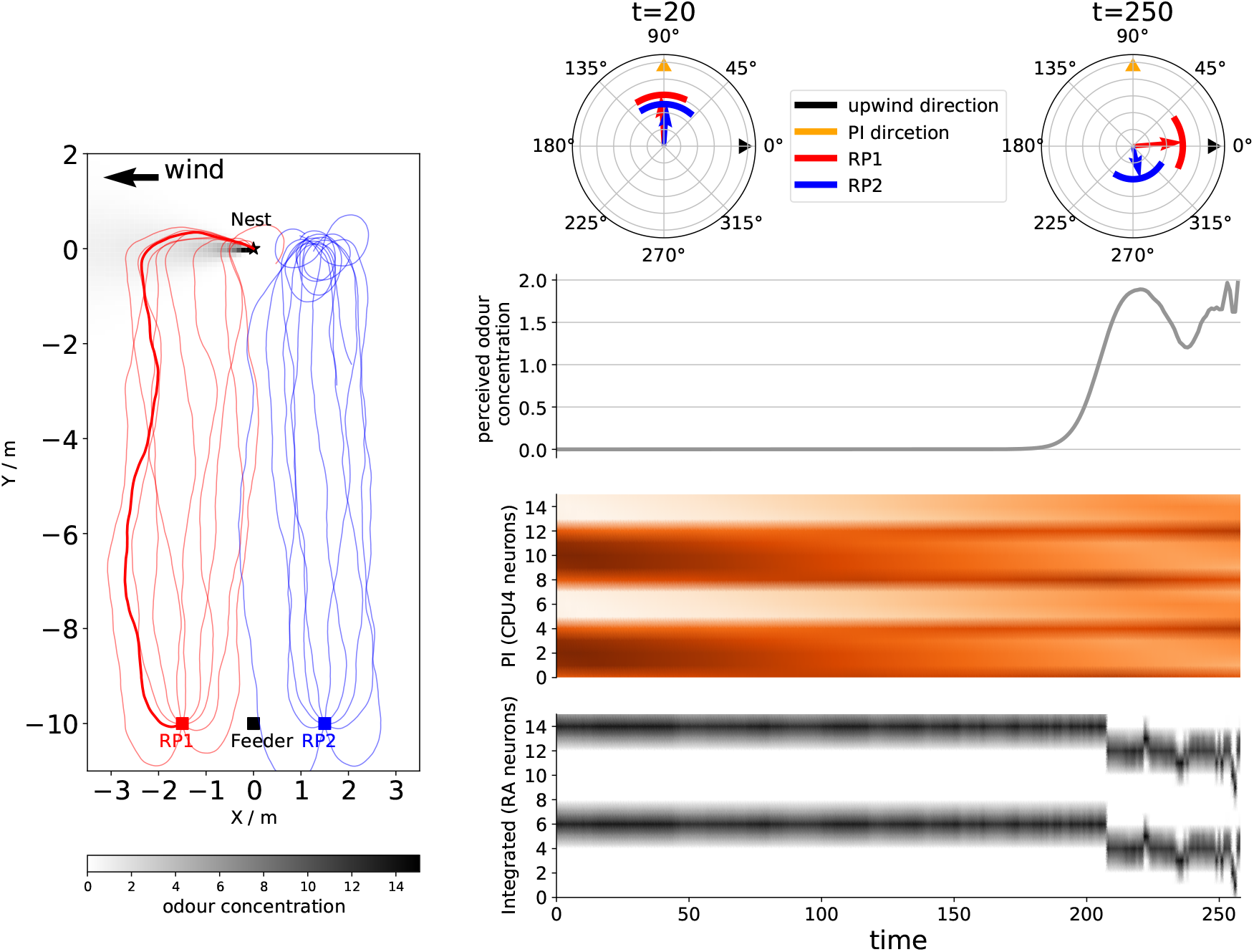
Left part draws the simulated ants’ homing paths and the group mean headings at *t* = 20 (when PI dominated) and *t* = 250 (when olfactory navigation should dominate the steering) are shown on the right. The instantaneous sensory value and neural activation of highlighted agent in the left panel during homing on shown on the right hand panel: from top to bottom, the value of perceived odour concentration, the activation of PI memory neurons (CPU4) and the ring attractor excitation neurons.

## Notes

### Competing Interest Statement

The authors have declared no competing interest.

